# Somatic symptom burden is associated with anger internalization measured as reduced reactive aggression

**DOI:** 10.64898/2026.11.09.765249

**Authors:** Ayano Arima, Satoshi Umeda

## Abstract

**Objectives:** Difficulties in emotion regulation, including anger suppression, have been linked to more severe somatic symptoms in patients with chronic pain. Because this evidence comes mainly from clinical samples, it remains unclear whether such difficulties precede or follow symptom chronification. We examined whether somatic symptom burden is associated with reduced reactive aggression, as a behavioral marker of anger internalization.

**Method:** Forty-four healthy adults (25 females) completed the Taylor Aggression Paradigm, a competitive reaction-time task in which they selected noise intensity delivered to a fictitious opponent across three provocation conditions (low, moderate, and high). somatic symptom burden was evaluated using the Somatic Symptom Scale-8 (SSS-8) and Patient Health Questionnaire-15 (PHQ-15). Alexithymia was evaluated using the Toronto Alexithymia Scale (TAS-20).

**Results:** Participants with higher somatic symptom burden selected lower noise intensities overall, with significant differences at low and moderate provocation levels. Difficulty identifying feelings tended to be positively associated with somatic symptom burden, whereas alexithymia was not associated with reactive aggression.

**Discussion:** These findings suggest that inhibited retaliation, previously reported in chronic pain patients, may already be present in healthy individuals with higher somatic symptom burden, potentially aiding early identification of chronic pain risk. Difficulty distinguishing emotions from bodily sensations may also contribute to somatic symptom expression.

## Introduction

Persistent physical symptoms are distressing bodily complaints that last for several months or longer, regardless of their cause. Because these symptoms occur in the context of somatic diseases, functional somatic disorders, mental disorders, and undiagnosed conditions, they are a central concern across a wide range of medical fields [1]. Headache, abdominal symptoms, and fatigue are among the most frequently reported symptoms in primary care, accounting for up to 50% of all consultations [2]. Somatic symptoms accumulate and persist over time, imposing a substantial burden on quality of life [3]. Regardless of its cause, physical distress, including pain (i.e., persistent physical symptoms), is not merely a sensory phenomenon but a complex experience that inherently encompasses emotional dimensions. How it is experienced varies substantially across individuals and within the same individual over time, depending on context, meaning, and psychological state [4]. Neuroimaging studies have revealed substantial overlap between the neural pathways underlying pain and emotion. In particular, the insula serves as a central hub for somatosensory processing and pain, and research on this region has highlighted the importance of internal bodily sensations in the subjective experience of emotion [5–7]. Given that pain and emotion share neural substrates and that somatic symptoms are closely intertwined with emotional processes, the way individuals regulate their emotions may play a key role in the development and maintenance of somatic symptoms. Therefore, understanding somatic symptoms from a psychological perspective is essential for the development and implementation of targeted interventions.

Emotion regulation (ER) refers to the processes by which individuals influence which emotions they have, when they have them, and how they experience and express those emotions [8]. Adaptive ER enables individuals to respond flexibly to stressors and maintain psychological homeostasis [9]. Difficulties with emotion regulation are associated with mental disorders and incorporated into several models of specific psychopathologies, including major depressive disorder, generalized anxiety disorder and social anxiety disorder [10,11]. Several reviews have recently examined the relationship between chronic pain and emotion regulation [12,13]. Schnabel et al. [14] found that patients with somatic symptom disorders and related conditions (SSD-RC) showed deficits in the identification stage of ER, particularly reduced emotional clarity and ER self-efficacy. SSD-RC were also significantly associated with deviant patterns of habitual strategy use (the selection stage), such as more frequent use of expressive suppression and less frequent use of cognitive reappraisal.

Among difficulties in emotion regulation, over-regulation has received particular attention for its association with somatic symptoms. Over-regulation refers to the consistent interruption of the natural resolution of emotional experiences through experiential avoidance [15] or expressive suppression [16]. Chronic reliance on over-regulation ultimately contributes to the formation and maintenance of somatic symptoms [17]. Previous studies have demonstrated that difficulties in emotion regulation increase the tendency to experience and express emotional distress as physical symptoms rather than processing it verbally or conceptually [18,19]. It has been conceptualized as somatization. Somatization is one psychological factor implicated in chronic pain. Lipowski [20] defined it as the tendency to experience and communicate somatic distress that is not explained by pathological findings, to attribute it to physical illness, and to seek medical help for it, typically in response to psychosocial stress. Rather than a disease or diagnostic category, somatization is an umbrella term for the experiential, cognitive, and behavioral characteristics of individuals who report physical symptoms despite limited medical findings [21]. Multiple psychological mechanisms, including cognitive processing, emotion regulation, and the perception of bodily sensations, contribute to the somatization process [21,22].

Anger offers a behavioral window into over-regulation: when anger is over-regulated, its suppression appears as reduced retaliation against provocation, which can be measured in the laboratory as reactive aggression. Reactive aggression is an impulsive, affective response to provocation that, unlike goal-directed proactive aggression, is closely tied to anger and positively associated with outward anger expression [23–26]; accordingly, ER strategies have been systematically linked to aggressive behavior [27]. In contrast, anger internalization refers to the tendency to experience anger while suppressing its outward expression [28]. In this pattern, aggressive impulses are inhibited and redirected internally through the accumulation of unexpressed emotion [29,30]. Previous studies have consistently linked anger internalization to somatization and somatic symptoms [31–33]. For example, patients with fibromyalgia tend to suppress their anger rather than express it [34,35]. Moreover, anger-suppression-induced sympathetic nervous system hyperactivation is associated with somatization [36]. These findings suggest that the maintenance of somatic symptoms and the suppression of aggressive behavior may arise through a shared mechanism of chronic over-regulation. However, these findings are primarily derived from studies of patients with chronic somatic symptoms.

Alexithymia is an individual difference factor that may underlie both somatization and the regulation of anger expression. It comprises difficulty identifying feelings and distinguishing them from bodily sensations (DIF), difficulty describing feelings (DDF), and externally oriented thinking (EOT) [37]. It’s conceptualized as a dimensional trait that varies along a continuum in the general population [38]. The concept was originally investigated as a basis of psychosomatic disorders [39]. Alexithymia is elevated in patients with chronic pain and associated with greater pain intensity and physical disability [40]. It is positively correlated with trait aggression and relational aggression [41,42], and difficulty identifying feelings is negatively correlated with anger control but positively correlated with aggression [43]. Therefore, alexithymia may influence reactive aggression through two opposing pathways. On the one hand, it may directly increase aggression. On the other hand, it may promote somatization and thereby indirectly reduce aggression.

However, these findings have two major limitations. First, most of them have been obtained from patients already diagnosed with chronic pain or somatic symptom disorders. It remains unclear whether anger internalization precedes the chronification of symptoms or develops as a consequence of living with chronic symptoms. Second, most previous studies have assessed anger suppression and emotion regulation using self-report measures [44,45]. However, individuals with a repressive coping style tend to underreport their negative emotions [46,47]. The stronger an individual’s tendency to suppress anger, the less likely self-report measures are to capture it. Indeed, few studies have experimentally examined emotion regulation in somatic symptoms [14].

This study examined whether somatic symptom burden, as an index of somatization tendency, is associated with anger internalization, as reflected in reduced reactive aggression. We assessed reactive aggression behaviorally with the Taylor Aggression Paradigm (TAP) [48]. The TAP is one of the most widely used laboratory measures of reactive aggression, and it allows researchers to manipulate provocation intensity in graded steps [49,50]. We had two hypotheses. First, we predicted that individuals with higher somatic symptom burden would show lower reactive aggression. Second, we predicted that alexithymia, as measured by the total score on the 20-item Toronto Alexithymia Scale (TAS-20), would be positively associated with somatic symptom burden. We also explored whether the association between somatic symptom burden and reactive aggression varies across provocation levels, how the DIF and DDF subscales of the TAS-20 relate to somatic symptom burden and reactive aggression, and whether DIF relates to reactive aggression indirectly through somatic symptom burden.

## Methods

### Study Design

This study followed a single-session laboratory design to examine the association between somatic symptom burden and reactive aggression, conducted in accordance with the Declaration of Helsinki and approved by the Keio University Research Ethics Committee (approval number: 230350000). Before participation, all individuals provided written informed consent, including consent for the use of collected data.

### Participants

A total of 44 healthy university students, graduate students, and working adults (17 males, 25 females, 2 non-responses; *Mean age = 21.9 years, SD = 1.58*) participated in this study, all of whom reported normal or corrected to normal vision. On the day of the experiment, participants verbally confirmed that they had not taken any medication affecting psychological or neurological functioning within three hours before the start of the session (including analgesics, cold medications, and antihistamines), had no history of cardiac disease, and had not consumed any vasoactive substances (including foods and beverages containing caffeine, alcohol, or nicotine).

### Procedure

#### Aggression Paradigm

The Taylor Aggression Paradigm (TAP) [48] is a competitive reaction-time task in which participants are led to believe they are playing against another participant. The general procedure was adapted from Böhnke et al. [51] and Rinnewitz et al. [52]. The task ran on a personal computer (HP EliteDesk 800 G6 SFF) and was programmed using PsychoPy; the task delivered auditory stimuli via headphones (Audio-Technica). To examine emotion regulation functioning in an interpersonal context, participants were led to believe that they were playing against a human opponent; following the experiment, they were debriefed that the opponent was, in fact, a computer program. Participants engaged in alternating competitive reaction-time trials against one of three fictitious opponents in separate rooms. Each opponent block comprised 15 consecutive trials, and the loser of each trial received an unpleasant noise stimulus from the opponent. Participants selected the intensity of the punishment that is, the volume of the noise on a scale of 1 to 10 on each trial. To standardize the aversiveness of the auditory stimuli, the experimenter played the noise file at its highest decibel level and gradually increased the PC volume until each participant found it intolerably aversive. The volume one level below this threshold then served as the maximum volume for the task. White noise served as the auditory stimulus, with 10 levels spaced at 1 dB, and the intensity delivered to the participant varied depending on the opponent. Each trial consisted of a decision phase and an outcome phase. During the decision phase, participants selected a punishment intensity from 1 to 10 using a selection bar centered on the screen, with 10 representing the highest intensity. A fixation cross then appeared for 1,000 ms, after which a red square appeared, and participants pressed the space key as quickly as possible. Reaction times outside the 150–450 ms window resulted in a loss; valid reaction times led to a win or loss determined with 50% probability. During the outcome phase, winners received notification that the noise at their selected intensity would reach the opponent. In contrast, losers received the noise at the opponent’s selected intensity, with visual feedback displaying the opponent’s chosen intensity. The task comprised three blocks of 15 trials each, with opponent provocation level varying across blocks: the opponent predominantly selected low intensities (levels 1–3) in the low provocation condition, moderate intensities (levels 4– 6) in the moderate provocation condition, and high intensities (levels 7–9) in the high provocation condition. Participants completed five practice trials before the main task. In the TAP, the mean punishment that participants deliver to their opponent (e.g., noise intensity and duration) is used as an index of reactive aggression. In the present study, participants selected only the intensity of the noise blast, with its duration held constant; therefore, the mean noise intensity delivered to the opponent was used as the measure of reactive aggression. Mean intensity was computed both for each provocation condition and across all conditions. Figure 1 illustrates the sequence of a single trial.

**Figure 1.**
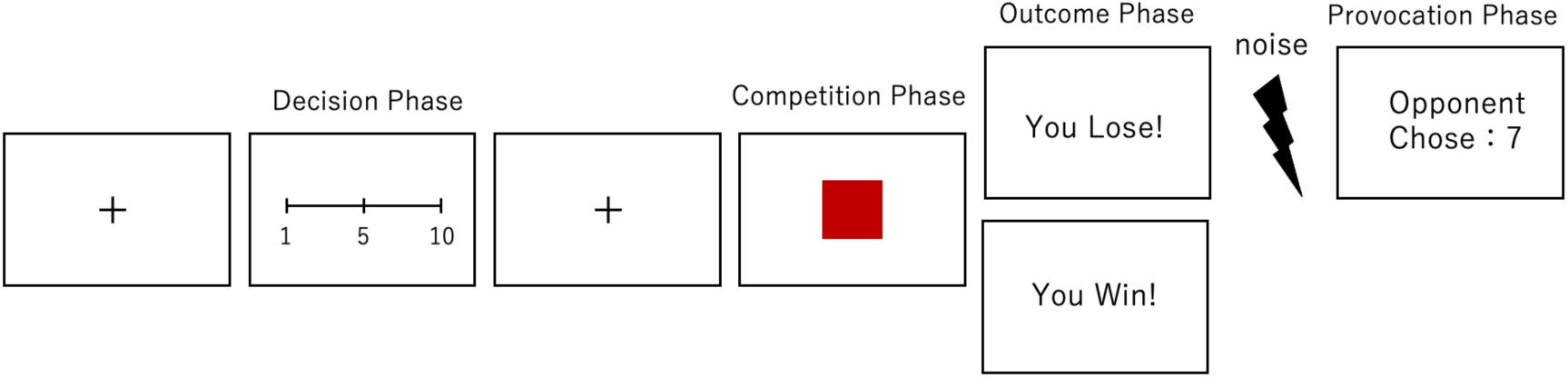
Trial sequence of the Taylor Aggression Paradigm. Each trial consisted of a decision phase and an outcome phase. Participants selected a noise intensity (1–10) to deliver to a fictitious opponent, then pressed the space key upon seeing a red square. Reaction times outside the 150–450 ms window resulted in a loss; valid reaction times led to a win or loss determined with 50% probability. Losers received noise at the opponent’s selected intensity. Three blocks of 15 trials varied in opponent provocation: low (1–3), moderate (4–6), and high (7–9).

#### Somatic Symptom Burden

Somatic symptom burden was assessed with the Japanese versions of the Somatic Symptom Scale-8 (SSS-8) [53,54] and the Patient Health Questionnaire-15 (PHQ-15) [55,56]. The SSS-8 is an eight-item questionnaire originally developed as an abbreviated version of the PHQ-15. Participants rated the degree of burden associated with eight somatic symptoms experienced over the past seven days on a five-point Likert scale (0 = not at all, 4 = very strong). In a German general population sample, the SSS-8 showed good reliability and validity, with psychometric properties comparable to those of the PHQ-15 [53]. For the PHQ-15, we used the Japanese symptom-assessment version [56], which corresponds to the Level 2 Somatic Symptom Severity measure recommended in the DSM-5 Cross-Cutting Symptom Measures [56,57]. Participants rated how much they had been bothered by 15 somatic symptoms over the past week on a three-point scale (0 = not bothered, 1 = bothered a little, 2 = bothered). Total scores range from 0 to 30, with scores of 0–4, 5–9, 10–14, and 15–30 indicating minimal, low, medium, and high symptom severity, respectively [55,56]. In the present sample, Cronbach’s α was .66 for the SSS-8 and .82 for the PHQ-15. Although the internal consistency of the SSS-8 was modest, its mean inter-item correlation (r = .19) fell within the range considered acceptable for measures of broad constructs [58], consistent with the scale’s coverage of heterogeneous symptom domains with a small number of items.

#### Alexithymia

The Toronto Alexithymia Scale (TAS-20) [59,60] is a 20-item self-report measure assessing three subdimensions of alexithymia: Difficulty Identifying Feelings (difficulty recognizing and distinguishing emotions and their associated bodily sensations), Difficulty Describing Feelings (difficulty verbally communicating emotions to others), and Externally Oriented Thinking (a tendency to focus on external facts). Each item is rated on a scale from 1 (strongly disagree) to 5 (strongly agree). Alexithymia is a personality trait originally proposed based on clinical observations of psychosomatic patients [39], and factor analytic studies have confirmed its three-factor structure comprising these subdimensions [61].

#### Group Classification and Statistical Analysis

Because the SSS-8 and PHQ-15 measure the same construct (r(42) = .89, p < .001), participants were assigned to groups only when both scales agreed. Those at or above the medium-severity threshold on both scales (SSS-8 ≥ 8 [53]; PHQ-15 ≥ 10 [55,56]) formed the higher somatic symptom burden group (n = 16), and those below it on both formed the lower group (n = 22). Six participants with discordant classifications (five meeting the threshold on the SSS-8 only and one on the PHQ-15 only) were excluded from group comparisons. Because of a technical error in data recording, TAP data were missing for one participant each in the low and moderate provocation conditions; analyses of mean TAP scores therefore included 42 participants. Group differences in TAP scores were examined using Welch’s t-tests, with Cohen’s d as the effect size. To examine the potential influence of sex, a supplementary linear regression including sex as a covariate was conducted for mean TAP scores. Associations among variables were examined using Pearson correlations, and p values for the exploratory correlations involving the DIF and DDF subscales were corrected using the false discovery rate (FDR) [62]. Exploratory simple mediation models (DIF → somatic symptom burden → mean TAP score) were tested using percentile bootstrap 95% confidence intervals (5,000 resamples). All analyses were conducted in R (version 4.6.1), using the lavaan package [63] for the mediation analyses, with two-tailed tests and α = .05.

## Results

### Descriptive Statistics

SSS-8 scores averaged 7.14 (SD = 4.71, range = 0–17). Based on the severity categories of Gierk et al. (2014), 11 participants were classified as having no to minimal (0–3), 12 low (4–7), 12 medium (8–11), 7 high (12–15), and 2 very high (16–32) somatic symptom burden. PHQ-15 scores averaged 7.80 (SD = 5.48, range = 0–18). The mean TAS-20 total score was 47.16 (SD = 10.7), comparable to the normative mean reported for non-clinical student samples (M = 47.40–48.99; Bagby et al., 1994). The mean DIF and DDF subscale scores were 15.70 (SD = 6.14) and 14.39 (SD = 5.03), respectively. Mean TAP scores were 4.03 (SD = 2.68) in the low provocation condition, 4.16 (SD = 2.23) in the moderate provocation condition, and 5.69 (SD = 2.81) in the high provocation condition. Descriptive statistics for the full sample and for the higher and lower somatic symptom burden groups are presented in Table 1.

**Table 1.** Descriptive Statistics for All Study Variables by Somatic Symptom Burden Group. Values are M (SD) for the full sample (N = 44) and for the higher (n = 16) and lower (n = 22) somatic symptom burden groups. The higher group met the medium-severity threshold on both the SSS-8 (≥ 8) and the PHQ-15 (≥ 10), and the lower group fell below it on both; six participants with discordant classifications are included only in the full sample. The number of participants varies across TAP conditions because of missing data. SSS-8 = Somatic Symptom Scale-8; PHQ-15 = Patient Health Questionnaire-15; TAS-20 = Toronto Alexithymia Scale-20; DIF = Difficulty Identifying Feelings; DDF = Difficulty Describing Feelings; EOT = Externally Oriented Thinking; TAP = Taylor Aggression Paradigm (mean noise intensity, 1–10).

| **Variable** | **All (N = 44)** | **Lower (n = 22)** | **Higher (n = 16)** |
|---|---|---|---|
| SSS-8 | 7.14 (4.71) | 3.27 (2.53) | 11.75 (2.65) |
| PHQ-15 | 7.80 (5.48) | 3.32 (2.51) | 13.81 (2.93) |
| TAS-20 | 47.16 (10.70) | 45.68 (10.35) | 49.69 (11.84) |
| DIF | 15.70 (6.14) | 14.36 (5.98) | 18.19 (5.97) |
| DDF | 14.39 (5.03) | 13.55 (5.31) | 15.19 (5.01) |
| EOT | 17.07 (3.57) | 17.77 (3.46) | 16.31 (3.75) |
| TAP: Low provocation | 4.03 (2.68) | 4.80 (3.15) | 2.99 (1.77) |
| TAP: Moderate provocation | 4.16 (2.23) | 4.86 (2.65) | 3.28 (1.50) |
| TAP: High provocation | 5.69 (2.81) | 6.40 (2.92) | 4.92 (2.72) |
| TAP: Mean | 4.56 (2.18) | 5.26 (2.36) | 3.73 (1.81) |

### Somatic Symptom Burden and Reactive Aggression

Consistent with our hypothesis, the higher somatic symptom burden group selected significantly lower noise intensities than the lower group in the low provocation condition (t(32.6) = 2.22, p = .033, d = 0.69) and the moderate provocation condition (t(32.6) = 2.30, p = .028, d = 0.71), as well as for the mean across all conditions (t(34.0) = 2.20, p = .034, d = 0.72; Figure 2). These effects were medium in size [64]. In the high provocation condition, the group difference was in the same direction but did not reach significance (t(33.7) = 1.61, p = .117, d = 0.52).

**Figure 2.**
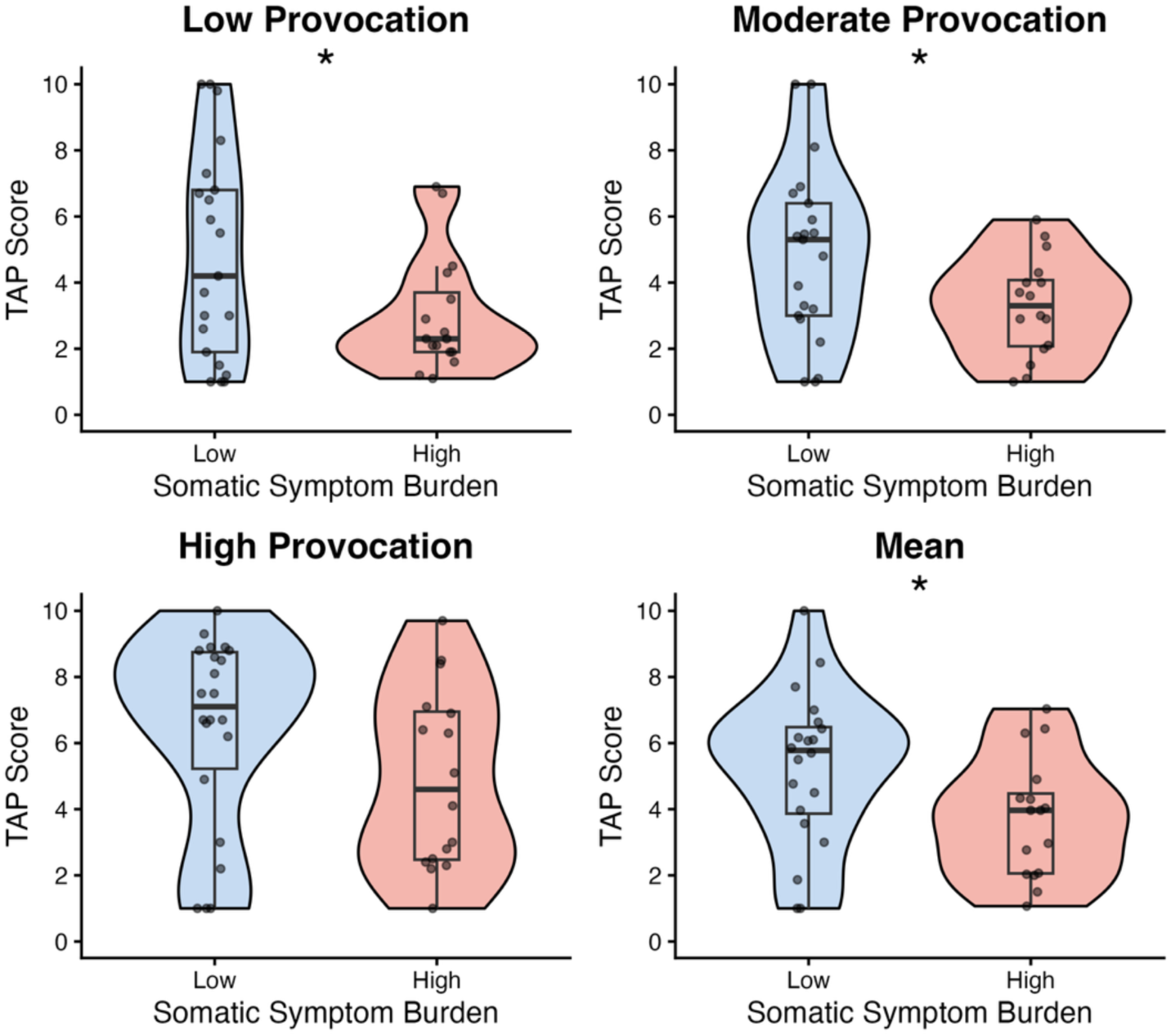
Comparison of Reactive Aggression Between the Higher and Lower Somatic Symptom Burden Groups. TAP (Taylor Aggression Paradigm) scores are the mean noise intensities (1–10) that participants selected for their opponent. The TAP score for each provocation condition is the mean within that condition. The TAP score for all conditions is the mean across the three conditions. Violin plots show the distribution of scores. Boxes indicate the median and interquartile range. Dots represent individual participants. The higher group (n = 16) met the medium-severity threshold on both the SSS-8 (≥ 8) and the PHQ-15 (≥ 10), and the lower group (n = 22) fell below it on both. *p < .05 (Welch’s t-test).

In a supplementary analysis, we examined the role of sex. The proportion of women was higher in the higher somatic symptom burden group (13 of 16, 81%) than in the lower group (7 of 19, 37%; Fisher’s exact test, *p* = .016). In a regression analysis including only participants who reported their sex (*n* = 35), the group difference in mean TAP scores (*B* = −1.76, *SE* = 0.69, *p* = .016) was reduced by approximately half and was no longer significant after controlling for sex (*B* = −0.94, *SE* = 0.71, *p* = .197). Female participants selected lower noise intensities than male participants (*B* = −1.85, *SE* = 0.72, *p* = .015).

### Correlations Among Somatic Symptom Burden, Alexithymia, and Reactive Aggression

When somatic symptom burden was treated as a continuous variable, SSS-8 scores were negatively but not significantly correlated with mean TAP scores (r(40) = −.24, 95% CI [−.50, .07], p = .133), whereas PHQ-15 scores were significantly negatively correlated with mean TAP scores (r(40) = −.32, p = .036). Regarding our second hypothesis, the TAS-20 total score was positively correlated with SSS-8 scores, as predicted, although this correlation fell just short of significance (r(42) = .29, 95% CI [−.00, .54], p = .054); its correlation with PHQ-15 scores was also positive but not significant (r(42) = .25, p = .099). The TAS-20 total score was not significantly correlated with mean TAP scores (r(40) = −.09, 95% CI [−.39, .22], p = .554).

### Associations of Alexithymia Subscales with Somatic Symptom Burden and Reactive Aggression

In exploratory analyses of the TAS-20 subscales, DIF was positively correlated with both SSS-8 scores (r(42) = .33, uncorrected p = .029) and PHQ-15 scores (r(42) = .35, uncorrected p = .022), but these correlations reached only marginal significance after FDR correction (both FDR-corrected *p* = .088). DDF was not significantly correlated with SSS-8 (r(42) = .26, FDR-corrected *p* = .170) or PHQ-15 scores (r(42) = .21, FDR-corrected *p* = .255). Neither DIF (r(40) = −.14, FDR-corrected *p* = .466) nor DDF (r(40) = .12, FDR-corrected *p* = .466) was significantly correlated with mean TAP scores.

### Somatic Symptom Burden as a Mediator Between Difficulty Identifying Feelings and Reactive Aggression

Because DIF showed a positive association with somatic symptom burden, we explored whether somatic symptom burden mediated the association between DIF and reactive aggression in the 42 participants with complete TAP data (Figure 3). When the PHQ-15 was the mediator, DIF was positively associated with PHQ-15 scores (a = 0.31, 95% CI [0.05, 0.58], β = .35), and PHQ-15 scores were negatively associated with TAP scores (b = −0.13, 95% CI [−0.22, −0.02], β = −.32). The indirect effect was significant, although the upper limit of its confidence interval was close to zero (ab = −0.039, 95% CI [−0.092, −0.001], standardized ab = −.11). Neither the direct effect (c’ = −0.01, 95% CI [−0.13, 0.11]) nor the total effect (c = −0.05, 95% CI [−0.17, 0.07]) of DIF on TAP scores was significant, indicating an indirect-only pattern. When the SSS-8 was the mediator, DIF was positively associated with SSS-8 scores (a = 0.25, 95% CI [0.03, 0.48], β = .34), but neither the association between SSS-8 and TAP scores (b = −0.10, 95% CI [−0.24, 0.03], β = −.21) nor the indirect effect (ab = −0.025, 95% CI [−0.075, 0.010], standardized ab = −.07) was significant. Given that the correlations between DIF and somatic symptom burden did not remain significant after correction, together with the cross-sectional design, the small sample, and the inconsistency between the two measures, these findings should be considered preliminary.

**Figure 3.**
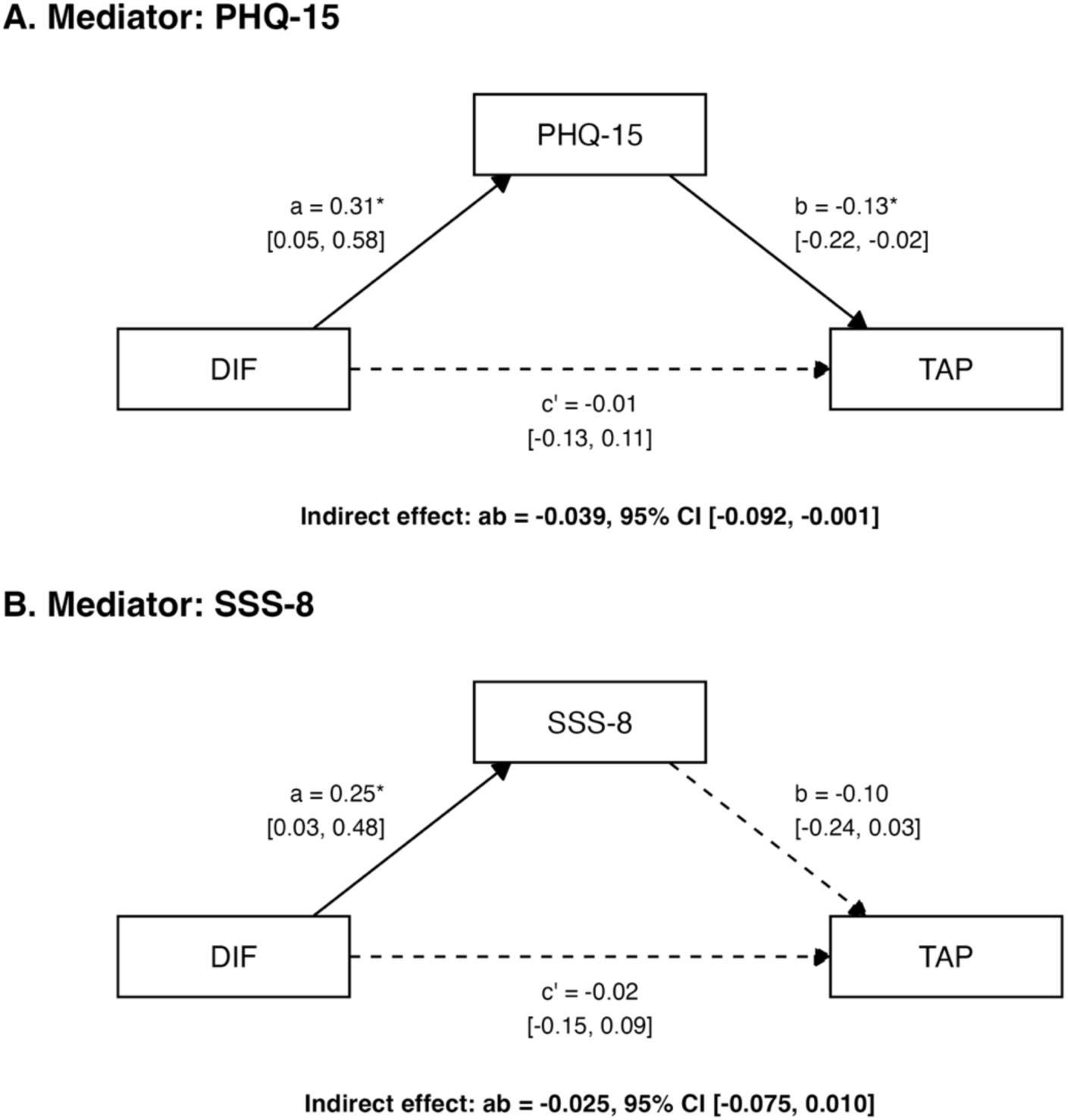
Exploratory Mediation Models of the Association Between Difficulty Identifying Feelings and Reactive Aggression. Both models tested whether difficulty identifying feelings (DIF), a subscale of the 20-item Toronto Alexithymia Scale (TAS-20), was indirectly related to reactive aggression through somatic symptom burden. Reactive aggression was indexed by the Taylor Aggression Paradigm (TAP) score for all conditions (mean noise intensity, 1–10). Panel A uses the Patient Health Questionnaire-15 (PHQ-15) as the mediator. Panel B uses the Somatic Symptom Scale-8 (SSS-8). Path a links DIF to the mediator, and path b links the mediator to TAP. The direct effect is denoted c’, and the indirect effect ab. Coefficients are unstandardized, with percentile bootstrap 95% confidence intervals in brackets (5,000 resamples, n = 42). Solid lines indicate paths whose 95% CIs did not include zero. Dashed lines indicate nonsignificant paths. *95% CI does not include zero.

## Discussion

This study aimed to examine whether somatic symptom burden is associated with reactive aggression, assessed behaviorally with the TAP, in healthy adults. Based on the anger internalization pattern reported in patients with fibromyalgia [34,35], we hypothesized that individuals with higher somatic symptom burden would show lower reactive aggression. The results were consistent with this hypothesis: participants with higher somatic symptom burden delivered less intense punishment to their opponents than those with lower burden, although this association was weaker when somatic symptom burden was treated as a continuous variable and was substantially attenuated when sex was taken into account. This finding extends previous evidence on anger internalization in chronic pain. A systematic review and meta-analysis of patients with chronic primary pain showed that most anger-related variables were significantly and positively correlated with pain and disability, with small-to-medium effect sizes, and suggested that, although the results were mixed, a greater tendency to hold anger in was associated with greater pain intensity and disability [65]. Moreover, an ecological momentary assessment study of patients with chronic low back pain showed that greater behavioral anger suppression at one time point predicted greater pain interference several hours later, even after controlling for anger at that time [44]. Aversive bodily states, including pain, have been linked to aggression and anger has been linked to chronic pain [66,67]. Against this background, the lower aggression among individuals with greater somatic symptom burden may be more consistent with the inhibition of the behavioral expression of anger than with a simple absence of anger. Aggressive behavior can serve as a means of escaping difficult emotional experiences and is closely linked to emotion regulation [29]. Therefore, individuals who habitually regulate negative emotions by inhibiting their outward expression may be less likely to retaliate even when provoked. This interpretation is also consistent with previous studies linking anger suppression to somatic symptoms [13,31,36,68]. Taken together, these findings suggest that the tendency to refrain from outward aggression, previously documented mainly in clinical samples, may also be observable in healthy individuals as an association with somatic symptom burden. However, inhibited anger may not simply dissipate but may instead be processed cognitively. According to the anger avoidance model, efforts to avoid anger can manifest as hostile rumination rather than aggressive behavior [69]. Moreover, anger rumination has recently been shown to be associated with somatic symptoms through negative affect [70]. Finally, the cross-sectional design leaves the direction of these associations open. For example, somatic symptoms may reduce the motivation or capacity to retaliate. Longitudinal studies are needed to determine whether anger internalization precedes symptom chronification.

DIF was positively correlated with both SSS-8 and PHQ-15 scores, although these correlations were only marginally significant after FDR correction. This pattern tentatively suggests that difficulty distinguishing emotions from bodily sensations may contribute to the expression of emotional distress as somatic symptoms, in line with previous findings that alexithymia mediates the relationship between pain intensity and psychological distress [71]. In contrast, neither the TAS-20 total score nor its subscales were significantly correlated with TAP scores. Previous studies have reported positive correlations between alexithymia and trait or relational aggression [41,42], and difficulty identifying emotions has been associated with poorer anger control and greater aggression [43]. The absence of such an association here may partly reflect the specificity of the TAP as a behavioral measure of reactive aggression. An exploratory mediation analysis offers another possible explanation: DIF was indirectly associated with lower TAP scores through higher PHQ-15 scores, whereas neither its direct nor its total effect was significant. If DIF relates to aggression through multiple pathways—one via somatic symptom burden associated with lower aggression, and others associated with greater aggression, as suggested by previous studies—these pathways may offset each other, resulting in no overall association. However, the indirect effect was small, and its confidence interval approached zero. It also did not reach significance when the SSS-8 served as the mediator, although the path from the SSS-8 to TAP scores was in the same direction. Similarly, PHQ-15 scores, but not SSS-8 scores, were significantly correlated with TAP scores. This discrepancy may reflect differences in symptom coverage between the two scales. Compared with the SSS-8, the PHQ-15 additionally assesses symptoms such as heart pounding or racing and fainting spells, and it assesses gastrointestinal symptoms in more detail. Many of these symptoms reflect autonomic arousal, which overlaps with the bodily components of emotion. Because DIF involves difficulty distinguishing emotions from such bodily sensations, the broader item set of the PHQ-15 may have captured aspects of somatization that the SSS-8 did not. Alternatively, the lower internal consistency of the SSS-8 relative to the PHQ-15 in this sample (α = .66 vs. .82) may have reduced its sensitivity. This finding should therefore be regarded as preliminary and tested in larger samples.

These findings likely capture only one facet of somatization, which also involves psychological factors beyond anger internalization [72]. Pain catastrophizing also serves as a key factor in amplifying pain experience and driving somatic symptoms [73,74], as negative cognitive appraisals promote excessive attention to bodily sensations, thereby maintaining and exacerbating symptoms. Interoception is also associated with somatization tendency. Interoception refers to the perception of the body’s internal states, a function that continuously represents the internal environment and operates at both conscious and unconscious levels [75]. Heightened responsiveness to signals from within the body may increase symptom reporting [76,77] though some studies find no differences in interoceptive accuracy between individuals with somatic symptom disorder and healthy controls, indicating that findings remain inconsistent [78]. Individual differences in interoception moderate both subjective and physiological indices of emotional experiences and regulation, and interoceptive awareness appears to play an important mediating role in self-assessed health status, particularly in the perception of pain [79]. The fear-avoidance model further proposes that pain-related fear, avoidance, and hypervigilance to bodily signals maintain somatic symptoms [80,81]. In future research, integrating measures of anger suppression, pain catastrophizing, interoceptive sensitivity, and fear-avoidance propensity to examine their relative contributions to the development of somatic symptoms remains an important challenge in elucidating the mechanisms of somatization.

### Limitations

This study has several limitations. First, we did not assess subjective emotional experiences during the TAP (Taylor Aggression Paradigm). The quality and intensity of negative emotions may vary considerably across individuals. Real-time ratings of emotional valence during the outcome phase would enable a more fine-grained examination of the emotional processes underlying aggression. Second, white noise served as the aversive stimulus, and we controlled its aversiveness through individual volume calibration. However, white noise is also used for mood regulation and pain relief [82], so some participants may not have perceived it as aversive. Future studies could use alternative or multimodal aversive stimuli to strengthen the ecological validity of the paradigm. Third, the pattern of reduced reactive aggression observed here may differ qualitatively from that in clinical populations such as patients with chronic pain. Directly comparing non-clinical and clinical samples would clarify how this pattern changes during the transition to symptom chronification. Finally, the replicability of the present findings should be confirmed under more robust conditions, such as larger and more diverse samples with a balanced sex distribution across groups. Such replication would provide more precise estimates of effect sizes and further clarify the association between somatization tendency and reactive aggression.

## Conclusion

Among the many psychological factors implicated in somatic symptoms, this study focused on anger internalization, a factor related to emotion regulation. We found that somatic symptom burden was associated with anger internalization measured as reduced reactive aggression, even in healthy individuals. This finding supports viewing somatic symptoms along a continuum that extends from healthy to clinical populations, rather than as a feature confined to somatic symptom disorders. From this dimensional perspective, anger internalization may be an emotional process involved in somatization. Future studies of this process could open avenues for the early identification of individuals at risk of symptom chronification and for preventive interventions.

## Funding

This work was supported by JSPS KAKENHI Grant Numbers JP22K18264, JP24H00177.

## Additional Information

The authors have declared no competing interest.

## Declaration of generative AI use

The authors used Claude (Anthropic; Claude Sonnet 4.6) to improve the English language of the manuscript. The authors have reviewed and edited the output and take full responsibility for the content of this publication.

## Data availability

The datasets generated and/or analysed during this study are not publicly available in order to protect participant confidentiality, but a de-identified dataset may be available from the corresponding author upon reasonable request.

## Author Contributions

A.A. and S.U. conceived and designed the study. A.A. developed the experimental task, collected the data, performed the statistical analyses, and wrote the original draft of the manuscript. S.U. supervised the project, acquired funding, and critically reviewed and revised the manuscript. Both authors approved the final version of the manuscript and agree to be accountable for all aspects of the work.

## Acknowledgments

We would like to thank Kazushi Shinagawa for technical support with the experimental program. We are also deeply grateful to all the participants of this study for their willingness to contribute their time and effort.

